# Developing an ethanol utilisation pathway based NADH regeneration system in *Escherichia coli*

**DOI:** 10.1101/2021.09.08.459383

**Authors:** Wenfa Ng

## Abstract

Many industrially relevant biotransformation in whole-cells are dependent on cofactors such as NADH or NADPH. Cofactor regeneration is an established approach for providing a cheap source of cofactors in support of the main biotransformation reaction in biocatalysis. In essence, cofactor regeneration uses a sacrificial substrate to help regenerate a cofactor consumed by the main biotransformation reaction. Enzymatic in nature, alternative cofactor regeneration systems with high efficiency and which utilises low cost sacrificial substrate are of interest. Glucose dehydrogenase system has been dominant in NADH regeneration. But, in its current incarnation, glucose dehydrogenase system is relatively inefficient in regenerating NADH with theoretical yield of one NADH per glucose molecule. This work sought to explore the utility of a two-gene ethanol utilisation pathway in NADH regeneration. Comprising the first step that takes ethanol to acetaldehyde, and a second step that converts acetaldehyde to acetyl-CoA, NADH from both steps could be mined for supporting biotransformation reaction in cofactor regeneration mode. Theoretically, ethanol utilisation pathway (EUP) affords a higher NADH yield of two NADH per ethanol molecule, and is therefore more efficient than glucose dehydrogenase (GDH) system. In this project, the EUP pathway was coupled to a cpsADH (an alcohol dehydrogenase from *Candida parapsilosis*) mediated ketone to alcohol anaerobic biotransformation with concentration of alcohol product as marker for efficiency of cofactor regeneration. Experiment tests showed that EUP was more efficient than GDH. Further, EUP could support biotransformation of both butanone and acetophenone in single and two-phase biotransformation, respectively. Additional work conducted to improve biotransformation efficiency revealed that ethanol provision positively correlated with biotransformation efficiency. Growing cell biotransformation was also found to improve biotransformation efficiency compared to resting cell due largely to the driving force generated by cell growth. Tests of a growth medium effect also found that cells cultivated in M9 ethanol medium delivered higher biotransformation efficiency compared to those cultivated in LB medium. This could arise due to the lower expression of NADH dependent enzymes during growth in M9 ethanol medium compared to LB medium that allowed more NADH to be diverted to support ketone biotransformation. However, a persistent problem with the experimental system is the relatively poor consumption of ethanol that points to need for further engineering of the system. Collectively, pathway-based NADH regeneration is possible with ethanol utilisation, with biotransformation efficiency dependent on mode of biotransformation (resting cell versus growing cell) and growth medium used.

## Introduction

Cofactor dependent oxidoreductases play a hugely important role in biocatalysis and biotransformation. Specifically, these enzymes are the pillars on which we could perform difficult organic chemistry kinetic resolution or biotransformation in whole cells under mild conditions compared to harsh conditions used in synthetic organic chemistry.^1,2^ Dependence on cofactors naturally meant that provision of expensive cofactors on a stoichiometric basis is necessary for maintaining high product yield. The alternative solution is that of using cofactor regeneration systems for cycling cofactors between their oxidized and reduced states to provide needed cofactors for a biotransformation.^3,4^ Principally, a cofactor regeneration system utilizes an enzyme to convert a sacrificial substrate to a second product that in the process also help regenerate the cofactor needed by the main biotransformation reaction.

As construed, the enzymatic cofactor regeneration system works in tandem with the main biotransformation reaction to help convert the main biotransformation substrate into the target product. Such a system would only work and remain coupled if the rates of the cofactor regeneration reaction is comparable to that of the main biotransformation reaction. But, the main benefit of a cofactor regeneration system is in shifting the cost burden from provision of expensive cofactor to that of the sacrificial substrate, which could be low-cost. Another less often noted concern in cofactor regeneration is the generation of a byproduct that could complicate the downstream separation of the main biotransformation product. Thus, work in the biocatalysis field has been on the search for low-cost sacrificial substrate/enzyme system that could be usefully tapped for cofactor regeneration without posing separation concerns for the main biotransformation product.

In the area of NADH regeneration, glucose dehydrogenase enzyme remains the most popular and dominant.^5-8^ Other NADH regeneration systems include formate dehydrogenase,^9-11^ and NADH oxidase.^12-14^ To be effective and useful, a cofactor regeneration system should deliver high yield of cofactor per molecule of substrate, and less often noted, be of relatively fast kinetics. For example, glucose dehydrogenase available in *Bacillus* species converts glucose to gluconolactone with the generation of one NADH.^15^ This meant a cofactor yield of one NADH per glucose molecule. Formate dehydrogenase, on the other hand, converts formate to carbon dioxide with the generation of one NADH. But, the system suffers from potential problem with the acidification of the medium due to carbon dioxide dissolution. Finally, NADH oxidase offers the most tantalizing prospects of a cofactor regeneration system delivering a benign product. Specifically, NADH oxidase uses molecular oxygen to convert NAD^+^ to NADH with the generation of either H_2_O_2_ or water.^16^ But, dependence on molecular oxygen meant that NADH oxidase could only operate under aerobic conditions, which restricts its application space in biocatalysis.

This work sought to introduce the NADH cofactor regeneration potential of a two-gene ethanol utilisation pathway that sequentially converts ethanol to acetaldehyde and on to the growth promoting metabolite, acetyl-CoA. The first step of the pathway is mediated by *adh2* from *Saccharomyces cerevisiae* with the generation of one NADH.^17^ This is followed by the second step of the pathway mediated by *ada* from *Dickeya zeae* that generates another NADH.^17^ Hence, in total, the ethanol utilisation pathway generates two NADH per ethanol molecule, and is more efficient in cofactor regeneration than the glucose dehydrogenase system. More importantly, the two-gene pathway ends in acetyl-CoA, which can be connected to central carbon metabolism, thereby, enabling distribution of ethanol flux to growth processes in growing cell biotransformation, that theoretically, could provide a greater driving force for NADH regeneration. In addition, acetyl-CoA remains in the cell and would not complicate downstream separation of biotransformation product. Finally, lack of oxygen dependence meant that the ethanol utilisation pathway (EUP) could operate under both aerobic and anaerobic biotransformation conditions, which expands the operating envelope of the cofactor regeneration system. Previously, ethanol utilisation pathway has been exploited for NADPH regeneration through a two gene pathway that converts ethanol to acetaldehyde and on to acetate. This project sought to convert ethanol to acetyl-CoA with NADH regeneration, that could be usefully tapped for biocatalysis.

In theory, cofactor regeneration could be implemented *in vitro* or *in vivo*. The latter is more attractive as whole-cell biocatalysis delivers better performance after issues such as enzyme cost and stability are taken into account. This study chose a whole cell biocatalytic system with microbial chassis as *Escherichia coli* to deliver a robust platform that could afford fair comparison of EUP and glucose dehydrogenase (GDH) capability at NADH regeneration. Given the objective of comparing the cofactor regeneration capability of a new cofactor regeneration system, the coupling biotransformation reaction is chosen as the ketone to alcohol biotransformation mediated by a proven NADH-dependent alcohol dehydrogenase (cpsADH) from *Candida parapsilosis*.^18^ Two substrates, acetophenone (hydrophobic) and butanone (hydrophilic) are chosen to test the utility of EUP in regenerating NADH under circumstances of a two-phase or single aqueous phase biotransformation system common in biocatalysis.

As a technology testbed, the *E. coli* BL21 (DE3) system transformed with the EUP pathway on one plasmid, and cpsADH on another plasmid was taken through a series of tests to demonstrate its utility in regenerating NADH in support of a ketone to alcohol biotransformation. After a straight comparison of cofactor regeneration capability between cells with EUP pathway and GDH system, the project moved to tune the parameters that theoretically could enable higher biotransformation efficiency to be obtained. This include looking into effects of ketone concentration and amount of ethanol provided. But, more important, a crucial test was whether growing cells could deliver higher biotransformation efficiency, which would validate theoretical postulations of how pathway that terminates in a growth promoting metabolite could deliver a driving force for substrate consumption, and, in this case, greater NADH regeneration. Resting cells refers to cells suspended in phosphate buffer without provision of nitrogen source. Finally, effect of growth medium on biotransformation efficiency may be transmitted by differing gene expression pattern, and this forms the basis for another segment of the research programme. In all these experiments, a key parameter to monitor is the amount of ethanol consumed. This is especially important given that a cofactor regeneration system could only be useful if there is no bottleneck that impedes substrate consumption and NADH regeneration. All experiments are conducted under anaerobic conditions to mimic industrial biocatalytic reactions without provision of oxygen through aerators in fermentation tanks.

Experimental results revealed that *E. coli* harbouring the reconstructed EUP pathway on a plasmid could grow on ethanol, which meant that the two-step EUP pathway is functional. More importantly, in biotransformation assays converting butanone to 2-butanol, product yield was enhanced with use of EUP for NADH regeneration relative to glucose/GDH system. This validated theoretical predictions that the EUP pathway is more efficient in regenerating NADH compared to glucose/GDH system. Furthermore, growing cells were shown to enable higher biotransformation efficiency compared to resting cells presumably due to the driving force generated by cell growth that push ethanol flux to acetyl-CoA. Finally, gene expression pattern resultant from cultivation in different growth media either prior to or during biotransformation also impact on biotransformation efficiency. In this case, M9 medium supplemented with 10 g/L ethanol was shown to enable higher biotransformation efficiency than LB medium in both resting cell and growing cell biotransformation.

The overall picture that emanates from the experimental results collected point to ethanol utilization as a viable platform for regenerating NADH in whole cell biocatalysis converting either acetophenone to 1-phenylethanol or butanone to 2-butanol in *E. coli*. Given that it is a plasmid-based platform, the two-gene EUP pathway could be readily transplanted to other species for enabling NADH regeneration in support of a NADH requiring biotransformation reaction after adjustment in plasmid replication origin and promoters to tailor to the genomic context of individual species. Although EUP is more efficient than the state-of-the-art glucose dehydrogenase system in regenerating NADH, higher cost of ethanol may still tilt the balance in favour of glucose dehydrogenase system. However, the scientific questions explored in this project should provide a firm foundation for future endeavours in tapping the NADH regeneration potential of EUP pathway such as in metabolic engineering applications where ethanol is the substrate providing the carbon source for building a target product molecule. In this case, downstream reactions whose enzymes require NADH would be able to use the NADH generated by EUP to achieve redox balance in the cell; thereby, ensuring a conducive cellular environment for attaining higher product yield and titer. Hence, ethanol/EUP system would then become a tool in the toolbox of metabolic engineers designing new pathways linking substrate to desired product where EUP provides a source of acetyl-CoA and NADH.

## Materials and Methods

### 1.4.1 Chemicals and medium composition

LB medium was purchased from Difco and used as is. Composition of LB medium was (g/L): Tryptone, 10.0; NaCl, 10.0; Yeast extract, 5.0. Composition of potassium phosphate buffer was (g/L): K_2_HPO_4_, 12.54; KH_2_PO_4_, 2.31. Composition of modified M9 medium was (g/L); K_2_HPO_4_, 6.8; KH_2_PO_4_, 3.0; NH_4_Cl, 1.0; NaCl; 2.0; Yeast extract, 1.0; Ethanol, 10.0. Composition of high cell density medium was (g/L): K_2_HPO_4_, 12.54; KH_2_PO_4_, 2.31; D-Glucose, 4.0; NH_4_Cl, 1.0; NaCl; 5.0; Yeast extract, 12.0; MgSO_4_, 0.24. Antibiotics used were ampicillin (100 µg/mL) and spectinomycin (50 µg/mL). Acetophenone, 1-Phenylethanol, glucose, butanone, 2-butanol, n-hexadecane and ethyl acetate were of analytical reagent grade (99%) and purchased from Sigma Aldrich.

### 1.4.2 Plasmid construction

Ethanol utilization pathway comprises two genes: *adh2* and *ada* were placed under the control of an autoinducible promoter, P_thrC3_ and inserted into a plasmid backbone with ampicillin as antibiotic resistance marker. Adh2 was cloned from *Saccharomyces cerevisiae* and ada was cloned from *Dickeya zeae* by my labmate, Ms Liang Hong, from whom I inherited the initial pathway. pMB1 serves as the origin of replication in this plasmid. On the other hand, cpsADH alcohol dehydrogenase enzyme was cloned from *Candida parapsilosis* and placed under the control of an autoinducible promoter, P_thrC3,_ and inserted into a plasmid backbone with spectinomycin antibiotic resistance marker. I inherited this plasmid from Mr. Liu Hongyuan from the lab. The replication origin of this plasmid is pA15. Given their differing antibiotic resistance marker and replication origin, both plasmids could be stably maintained in *E. coli* BL21 (DE3). Both plasmids were transformed into *Escherichia coli* BL21 (DE3) using standard heat shock transformation method.

### 1.4.3 General procedure for cloning of genes from bacterial species

Glucose dehydrogenase (gdh, *Bacillus subtilis*), aldehyde dehydrogenase (aldA, *E. coli*) and alcohol/aldehyde dehydrogenase (adhE, *E. coli*) were cloned from their respective bacterial species. Using appropriate primers, the respective genes were cloned through colony PCR in the thermocycler (Biorad, T-100). Specifically, 1 µL of a 16 hour bacterial culture in LB Miller medium was used as template for PCR amplification. PCR cycling conditions were: 98 °C for 5 mins, followed by 34 cycles of (98 °C for 8 sec, 55 °C for 15 sec, and 72 °C for 120 sec), with final extension of 72 °C for 5 mins. The PCR product was clean-up through gel electrophoresis (130V for 30 mins, Biorad power pack) followed by gel purification using Thermo Fisher Scientific gel purification kit (GeneJet Gel Extraction Kit, Cat No: #0692). The resulting gene fragment was ligated with the plasmid backbone using a ligase-free ligation method developed in the lab that includes incubation with 10 mM MgCl_2_ in a thermocycling step in a PCR thermocycler (Biorad, T-100). The cycling programme was as follows: 80 °C for 1 min, 68 °C for 10 mins, and infinite hold at 4 °C. DNA assembly method used was developed in the lab and is known as the GT standard.^19^

After plasmid ligation, colony PCR with appropriate primers were used to confirm ligation of gene fragment with plasmid backbone. Amplified gene fragment was extracted and purified with a gel purification kit (GeneJet Gel Extraction Kit, Cat No: #0692). Sequence of the gene fragment was verified by Sanger sequencing (BioBasic Singapore). Primers used for the experiments are as listed in Supplementary Table 1 below.

### 1.4.4 Cell cultivation

*E. coli* was inoculated in 10 mL LB medium with 100 µg/mL ampicillin and 50 µg/mL spectinomycin in 125 mL shake flask. Incubation conditions were 37 °C and 225 rpm for all cultivations. The incubator used was N-Biotek NB-205 Incubator-shaker. After 5 hours of cultivation, 0.25 mL of seed culture served as inoculum for 25 mL of LB medium with antibiotics in a 125 mL shake flask and the experiment cultures were cultivated for 20 hours. Optical density of the culture broth was measured at 600 nm with a UV-Visible spectrophotometer (Ambersham Biosciences). Appropriate dilution with deionized water was performed for samples with absorbance that exceed 1. Cells were harvested for biotransformation by centrifugation at 3300g for 5 minutes. Subsequently, cells were resuspended with 89 mM potassium phosphate buffer for a washing step, which included centrifugation at 3300g for 5 minutes. Cells were finally resuspended in 89 mM potassium phosphate buffer with a factor of 2 concentration in optical density for resting cell biotransformation. In growing cells biotransformation, cells were not washed with potassium phosphate buffer but were transferred into either LB Miller medium or M9 ethanol medium with a factor of 0.1 in optical density.

### 1.4.5 Acetophenone biotransformation

Given the poor solubility of acetophenone and 1-Phenylethanol in water, a two-phase system comprising n-hexadecane and potassium phosphate buffer was used in biotransformation. Specifically, either 5 or 20 g/L acetophenone was dissolved in 0.5 mL of n-hexadecane, while 0.5 mL of potassium phosphate buffer contained *E. coli* cells and 10 g/L ethanol. The reaction mixture was contained in 2.0 mL HPLC vial and incubated anaerobically at 37 °C and 225 rpm for 20 hours. Duplicate experiments were carried out. Cells were precultivated in LB medium for resting cells experiments.

At suitable time-points, HPLC vials were removed from the incubator. The contents were transferred to a 1.5 mL microcentrifuge tube and centrifuged at 20000g for 2 minutes. The upper organic phase containing the substrate and product was aliquoted and diluted with ethyl acetate prior to gas chromatography-mass spectrometry (GC-MS) analysis for detecting acetophenone and 1-Phenylethanol. Following that, the organic phase was carefully removed by pipetting; thereby, leaving behind an aqueous phase where 200 µL was aliquoted, filtered through a 0.22 µm nylon filter prior to high performance liquid chromatography (HPLC) analysis for detecting ethanol and acetate.

The performance measure selected for this segment of the work is yield per unit optical density, and is as defined below:

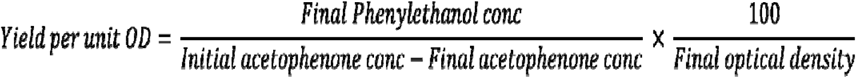

### 1.4.6 Butanone biotransformation

Since butanone and 2-butanol are highly water soluble, only a single-phase biotransformation system is required. Specifically, 10 g/L of butanone is added to either potassium phosphate buffer (resting cells) or growth medium (growing cells) together with 10 g/L of glucose or ethanol for biotransformation. Concentrations of butanone and ethanol were titrated in experiments aimed at understanding the effect of substrate toxicity effect or whether provision of higher ethanol concentration would drive higher biotransformation efficiency, respectively. The reaction mixture was contained in 2.0 mL HPLC vial and incubated anaerobically at 37 °C and 225 rpm for 44 hours. Triplicate experiments were carried out.

At suitable time-points, HPLC vials were removed from the incubator. The contents were transferred to a 1.5 mL microcentrifuge tube and centrifuged at 20000g for 2 minutes. 200 µL of the supernatant was aliquoted, filtered through a 0.22 µm nylon filter prior to high performance liquid chromatography (HPLC) analysis for detecting, butanone, butanol, ethanol or glucose and acetate.

The performance measure selected for this segment of the work is yield per unit optical density, and is as defined below:

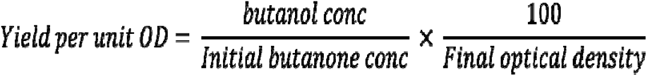

### 1.4.7 Gas chromatography mass spectrometry and high performance liquid chromatography analysis

Gas chromatography mass spectrometry was used in analysing the concentration of acetophenone and 1-Phenylethanol in Agilent 7890 GC and Agilent 5977B MSD. Injection volume used was 1 µL and the oven temperature profile used was 50 °C for 1 min, ramp from 50 to 180 °C at 10 °C/min, followed by another ramp from 180 to 280 °C at 50 °C/min and a hold at 280 °C for 3 minutes. MS detector was turned off at 13 minutes to protect the mass spectrometry detector.

High performance liquid chromatography was conducted with an Agilent 1260 series instrument equipped with a Bio-rad Aminex HPX-87H column. 5 µL of sample was injected into the column and eluted with 5 mM sulphuric acid at 0.7 mL/min. Isocratic elution was used and the column was maintained at 50 °C. Analytes were detected by a refractive index detector.

## Results and Discussion

### Integration of ethanol utilisation pathway with cpsADH alcohol dehydrogenase in a whole-cell biocatalytic system

At the theoretical level, the two gene ethanol utilisation pathway (EUP) delivers one NADH per step of the pathway that culminates in two NADH generated per ethanol molecule. This is a 100% improvement over the competing glucose dehydrogenase (GDH) system which regenerates one NADH per glucose molecule (Figure 1a). Hence, theoretically, the EUP pathway is more efficient than glucose dehydrogenase in regenerating NADH.

**Figure 1.**
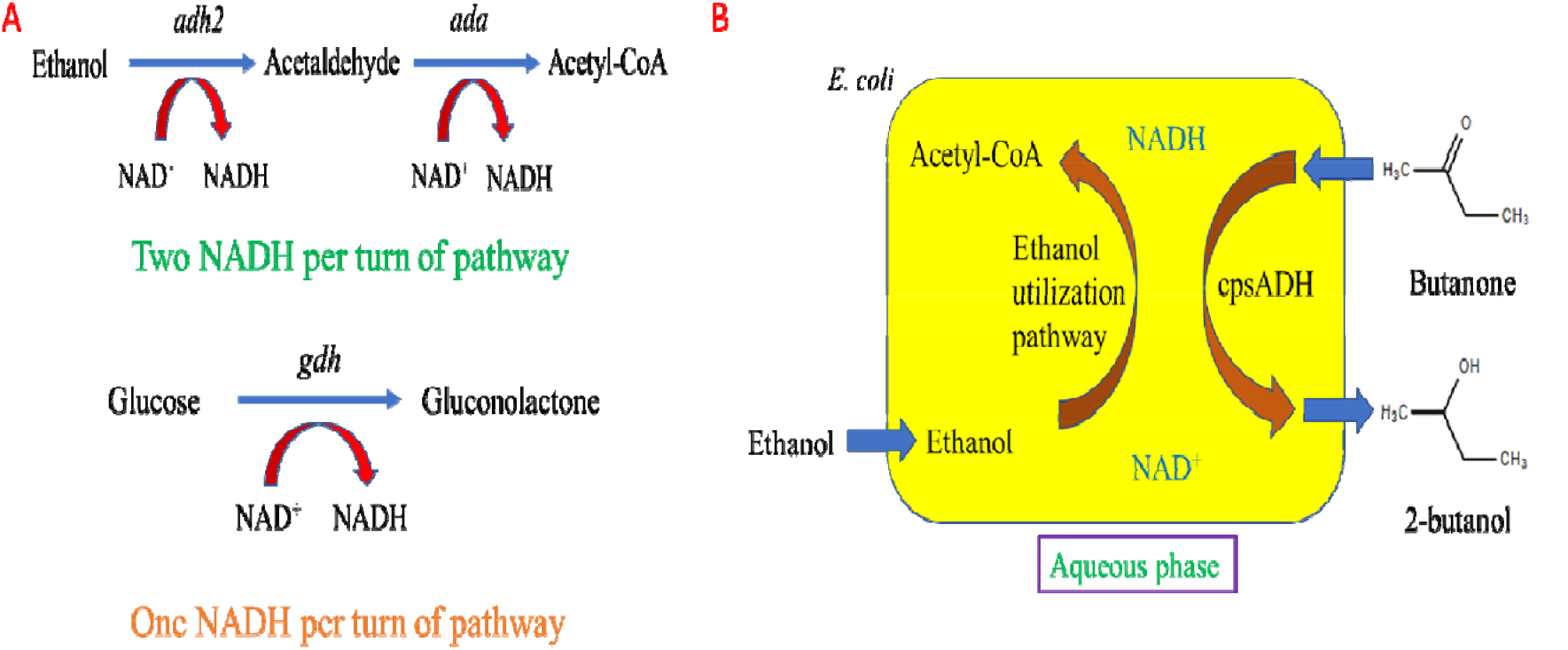
Conceptual basis of ethanol utilisation pathway (EUP) and whole cell biocatalysis. A) Ethanol utilisation pathway delivers higher NADH regeneration capability compared to glucose dehydrogenase system, B) Integration of EUP with cpsADH in an *Escherichia coli* BL21 (DE3) whole cell biocatalytic system.

To be effective, the EUP pathway must be integrated with cpsADH in a whole cell biocatalytic system with *E. coli* BL21 (DE3) as microbial chassis. Specifically, the two gene EUP pathway is encoded on a plasmid with pthrC3 autoinducible promoter and pMB1 replication origin, while cpsADH enzyme is encoded on another plasmid with pthrC3 autoinducible promoter and p15A replication origin. The whole cell biocatalytic system works by uptaking ethanol and converting it into acetyl-CoA with the generation of two NADH per ethanol molecule. At the same time, butanone or acetophenone would be taken up by the cell and converted by the NADH dependent cpsADH into alcohol which is secreted out of the cell (Figure 1b). Amount of 2-butanol or 1-phenylethanol produced would be taken as yardstick to assess the biotransformation efficiency of the cofactor regeneration system.

### Ethanol utilisation pathway could deliver growth to *E. coli* under aerobic and anaerobic conditions

Ethanol utilisation pathway ends with acetyl-CoA, which is a growth promoting metabolite. Thus, if the pathway is functional, provision of ethanol in M9 medium should engender growth. Experimental results in M9 + 10 g/L ethanol medium reveals that the EUP pathway was functional and could support growth of *E. coli* ethcps2 under aerobic conditions. Specifically, rapid growth of *E. coli* ethcps2 during exponential phase coincided with ethanol utilisation (Supplementary figure S1a). Similarly, *E. coli* ethcps2 could attain appreciable growth in M9 ethanol medium under anaerobic conditions at 37 °C and 225 rpm. This is consistent with theoretical predictions that the EUP pathway genes would be expressed under anaerobic conditions and that the enzymes are not dependent on oxygen for function. More significantly, *E. coli* ethcps2 could attain comparable growth in M9 medium supplemented with either 10 g/L glucose or ethanol (Supplementary figure S1b).

### EUP could regenerate NADH in support of acetophenone biotransformation in two-phase system

The first part of the project concerns the integration of the EUP cofactor regeneration system with cpsADH enzyme as a whole-cell biocatalytic system suited for the biocatalytic conversion of ketone to alcohol. Acetophenone, a hydrophobic substrate, would be the first substrate to be tested. To be useful, a promoter needs to drive the transcription and expression of EUP genes. In this project, a synthetic T7 promoter commonly used in biotechnology and an autoinducible pthrC3 promoter from *E. coli* are evaluated. As this was a resting cell experiment, *E. coli* cells harbouring EUP plasmid with T7 promoter were induced with 0.1 mM IPTG 3 hrs into cell cultivation. Experimental results revealed that biotransformation efficiency of acetophenone to 1-phenylethanol biotransformation was higher when EUP genes were under the control of pthrC3 compared to T7 promoter (Figure 2a). This came about presumably due to the higher expression of genes under the control of autoinducible pthrC3 promoter compared to T7 promoter.

**Figure 2.**
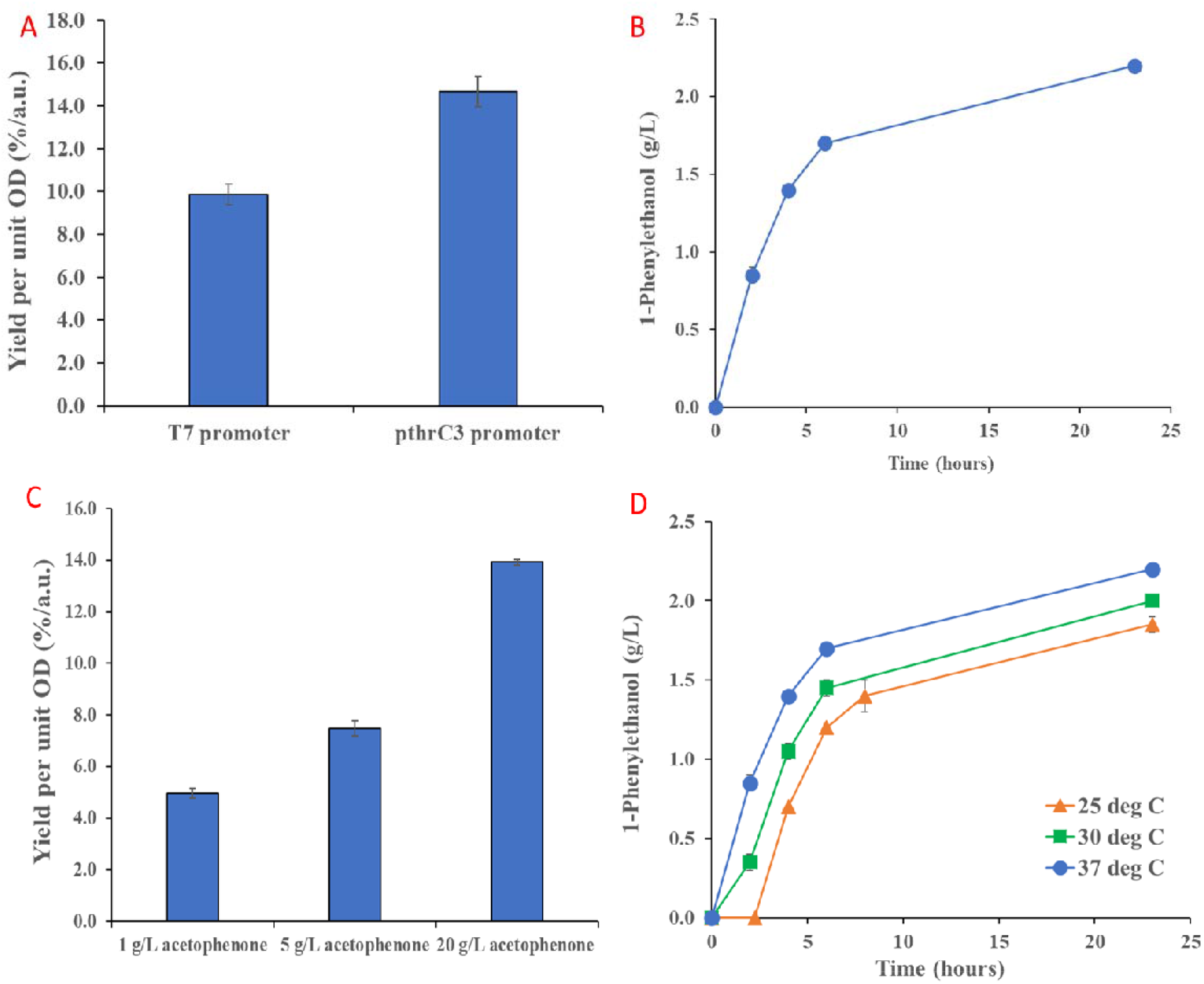
EUP could support acetophenone biotransformation in *E. coli* in a two phase system. A) Autoinducible pthrC3 promoter delivered higher biotransformation efficiency compared to T7 promoter in *E. coli* microbial chassis at 37 °C and 225 rpm in resting cells anaerobic biotransformation, B) Formation of 1-Phenylethanol product was rapid in the first few hours of the biotransformation, but tapered towards the latter stage of the process, C) Yield per unit OD improved with increase in acetophenone substrate concentration, thereby, indicating that the n-hexadecane organic phase could store and ameliorate potential toxicity effect from elevated acetophenone concentration, D) Higher incubation temperature favoured the formation of more 1-Phenylethanol product and greater biotransformation efficiency.

Kinetics of biotransformation is a hugely important parameter that governs the industrial feasibility of a given reaction. Experimental data in this area showed that the acetophenone to 1-phenylethanol biotransformation was quite rapid in the first few hours, but reaction slowed towards latter part of the reaction (Figure 2b). This points to possible equilibrium control of the biotransformation reaction as cpsADH is a reversible enzyme able to catalyze both directions of the biotransformation reaction. In any biotransformation system, desire is always to increase substrate loading to help improve product yield. Thus, acetophenone substrate loading experiments were carried out to determine the extent in which the approach would help push the equilibrium position of the reversible biotransformation experiment forward towards product formation. Data revealed that higher biotransformation efficiency was obtained with higher acetophenone loading moving from 1 g/L to 20 g/L acetophenone (Figure 2c). It is important to note that there was no substrate toxicity effect from the elevated concentration of acetophenone used. One possibility is that the organic n-hexadecane phase served as a reservoir for acetophenone, and this only allows small amount (solubility limit) of the substrate to come into contact with cells, thereby, helping to ameliorate toxicity effect of the substrate. In the current climate of reducing carbon footprint of biological processes, incubation temperature of biotransformation is an important parameter for tuning. With *E. coli* BL21 (DE3) as chassis, the typical process temperature would be 37 °C. This segment of the work sought to examine the possibility of reducing the process temperature to either 30 or 25 °C to help reduce heating requirement of the biotransformation. Results showed that this was not possible. In particular, product yield and biotransformation efficiency improved in a monolithic fashion with incubation temperature moving from 25 to 37 °C. This could likely be due to higher temperature enabling greater enzymatic activity and productivities.

### EUP cofactor regeneration supports butanone biotransformation in single phase biocatalysis

After verifying that the EUP pathway could regenerate NADH cofactor in support of a biotransformation reaction in two-phase system, work subsequently moved to determine the efficacy in which the EUP pathway could support a ketone to alcohol biotransformation reaction in single aqueous phase. Butanone is the substrate chosen due to its hydrophilic nature, and where the corresponding product, 2-butanol is a useful biofuel. Resting cells equipped with both EUP and cpsADH (i.e., *E. coli* ethcps2) were cultivated in M9 + 10 g/L ethanol medium for the resting cells experiment. A control experiment was also conducted where *E. coli* without EUP but with cpsADH (i.e., *E. coli* cps2) was taken through the same biotransformation protocol to examine the ability of the endogenous ethanol and glucose utilisation pathways in regenerating NADH. In all experiments, glucose was the reference substrate to serve as positive control as provide comparison for the relative utility of EUP and endogenous ethanol utilisation system in regenerating NADH.

Experiment results revealed that EUP could support butanone biotransformation to 2-butanol through regeneration of NADH. For comparison, the yield per unit OD of cells with EUP versus those with endogenous ethanol utilisation system was about 2 times (Figure 3a). More importantly, endogenous ethanol and glucose utilisation systems delivered similar biotransformation efficiency. Overall, the results suggest that the EUP pathway was functional and could support a whole cell biocatalytic reaction converting butanone to 2-butanol. Analyte concentration obtained by HPLC analysis of the single aqueous phase revealed that cells fed with ethanol also had the highest butanone uptake and utilisation (Figure 3b). This suggests a correlation between NADH regeneration and butanone consumption, which strongly implies that NADH availability drives butanone uptake. To be a useful cofactor regeneration system, EUP must enable high consumption of ethanol to regenerate as much NADH as possible in support of biotransformation reaction. But, experimental profile of ethanol utilisation efficiency revealed a large residual amount of ethanol in the reaction system even after 2 days of incubation with only 2-3 g/L of ethanol utilized (Figure 3c). This indicates that there is much room for further driving ethanol utilisation and cofactor regeneration for delivering higher product yield. At the same time, the data also suggests a possible fundamental constraint in ethanol utilisation that could likely be associated with anaerobic conditions used in the biotransformation. The EUP pathway ends in acetyl-CoA. For resting cells, acetyl-CoA could accumulate as there is a lack of growth processes that siphon off the produced acetyl-CoA. But, experimental data suggests that acetate may be the final product of EUP operation in resting cells given its relatively high titer and progressive increase in concentration with incubation time (Figure 3d). What likely transpires in resting cells fed with ethanol and undergoing biotransformation is that acetyl-CoA from EUP was transformed to acetate through acetyl-phosphate.

**Figure 3.**
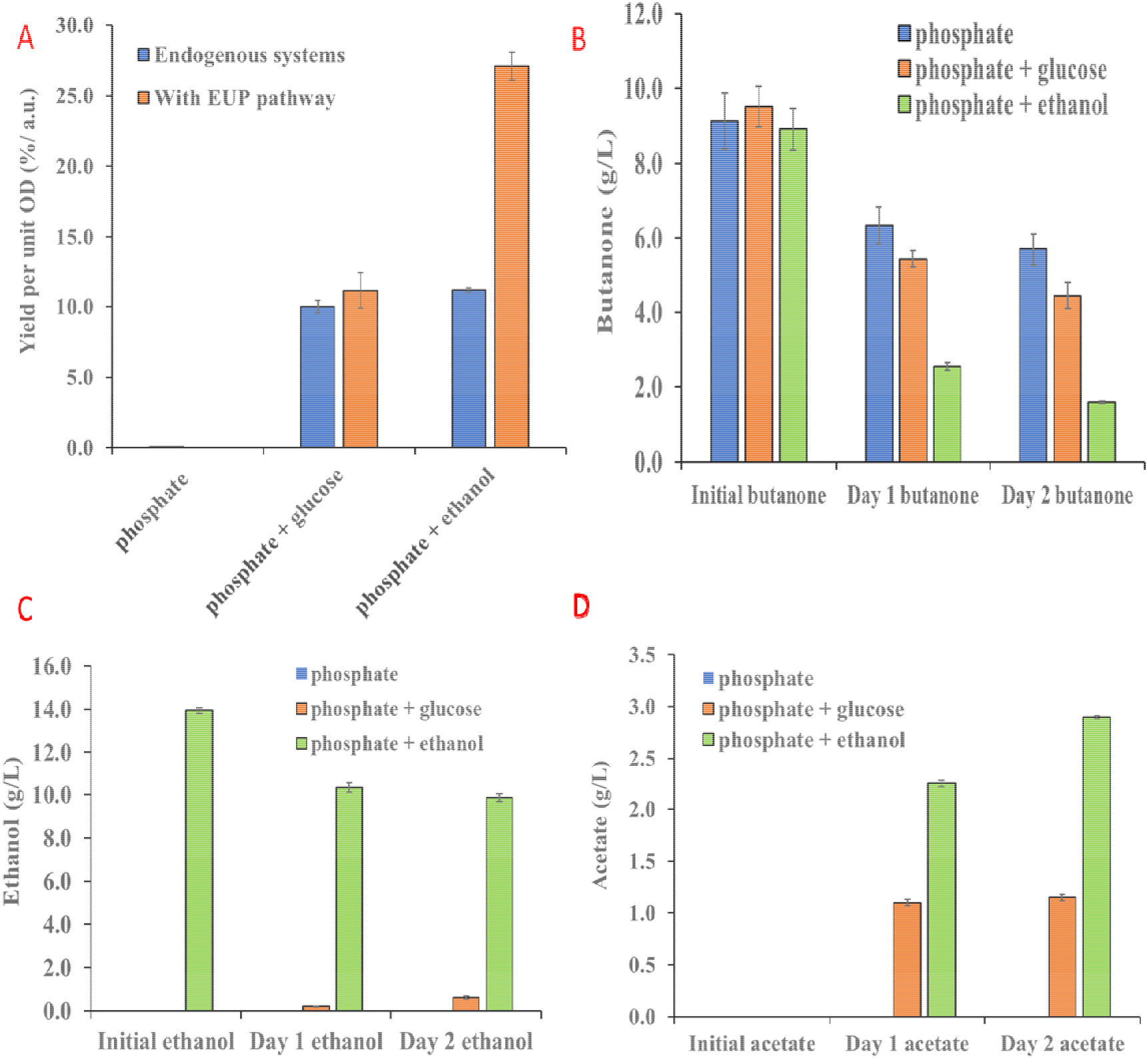
EUP enabled butanone biotransformation in single phase system in resting cells precultivated in M9 + 10 g/L ethanol medium at 37 °C and 225 rpm. A) EUP pathway delivered higher biotransformation efficiency compared to endogenous glucose and ethanol metabolization systems, B) Butanone consumption was highest in cells fed with ethanol, thereby, indicating a correlation between NADH regeneration and butanone utilisation, C) Ethanol utilisation was poor over the two day incubation period, which points to room for engineering higher ethanol utilisation, D) Acetate formation was highest in cells fed with ethanol, which indicates that acetyl-CoA conversion to acetate is a major route for channelling ethanol pathway flux in resting cells.

Efforts were also directed towards tuning parameters such as ethanol and butanone concentration to improve biotransformation efficiency. This arises because both the EUP pathway and cpsADH power reversible reactions. To improve NADH regeneration and product formation, ethanol and butanone concentration would need to be increased to push the equilibrium position forward towards product formation. In terms of ethanol provision, experiment results elucidated a positive correlation between ethanol concentration and biotransformation efficiency for resting cells (Supplementary Figure S2a). This indicates that higher ethanol concentration pushes flux towards acetyl-CoA in EUP and results in higher regeneration of NADH. However, such positive correlation was not observed in growing cell biotransformation where 10 g/L ethanol exerted some cellular toxicity. On the other hand, high substrate loading is always desired in biotransformation reactions to improve product titer. In this area, data revealed a clear substrate toxicity effect from single phase biotransformation (Supplementary Figure S2b). Specifically, higher butanone concentration resulted in a decline in biotransformation efficiency for resting cells precultivated in either LB or M9 + 10 g/L ethanol medium. Such toxicity effect could arise from the elevated butanone concentration experienced by the cells during biotransformation.

### Comparison of biotransformation efficiency and NADH regeneration between EUP and GDH

Prior experiments have demonstrated the utility of EUP in regenerating NADH in support of a ketone to alcohol biotransformation occurring in *E. coli* BL21 (DE3) whole cells. But, we do not yet know its relative efficacy in regenerating NADH and promoting biotransformation compared to other systems such as glucose dehydrogenase (GDH). Thus, work started in this project to clone GDH from *Bacillus subtilis* to serve as comparison for the EUP system. GDH gene was successfully cloned by PCR and inserted into the same vector as EUP under the control of the same autoinducible pthrC3 promoter to afford fair comparison between the two NADH regeneration systems. Similar to EUP, the plasmid encoding GDH was co-transformed with plasmid encoding cpsADH into *E. coli* BL21 (DE3) to constitute strain *E. coli* GDHcps1 for biotransformation.

In resting cells experiment with *E. coli* cps2 fed with either 10 g/L glucose or 10 g/L ethanol as controls, EUP pathway delivered higher biotransformation efficiency compared to GDH irrespective of whether LB or M9 + 10 g/L ethanol was used for precultivation (Figure 4a). Such data corroborate theoretical reasonings that EUP would deliver higher NADH regeneration efficiency compared to GDH as EUP regenerates two NADH molecule per ethanol molecule compared to GDH regeneration of one NADH per glucose molecule. At the same time, both GDH and EUP delivered higher biotransformation efficiency compared to their respective endogenous glucose or ethanol utilisation systems, which indicated that heterologous expression of GDH and EUP were successful and both cofactor regeneration systems were functional. Butanone utilisation per unit OD also provided useful yardsticks for assessing the relative performance of GDH and EUP. Specifically, data revealed that butanone utilisation was higher for cells with EUP compared to those with GDH (Figure 4b). This suggests that NADH regeneration efficiency is intimately tied to butanone consumption, which higher NADH regeneration driving higher butanone consumption.

**Figure 4.**
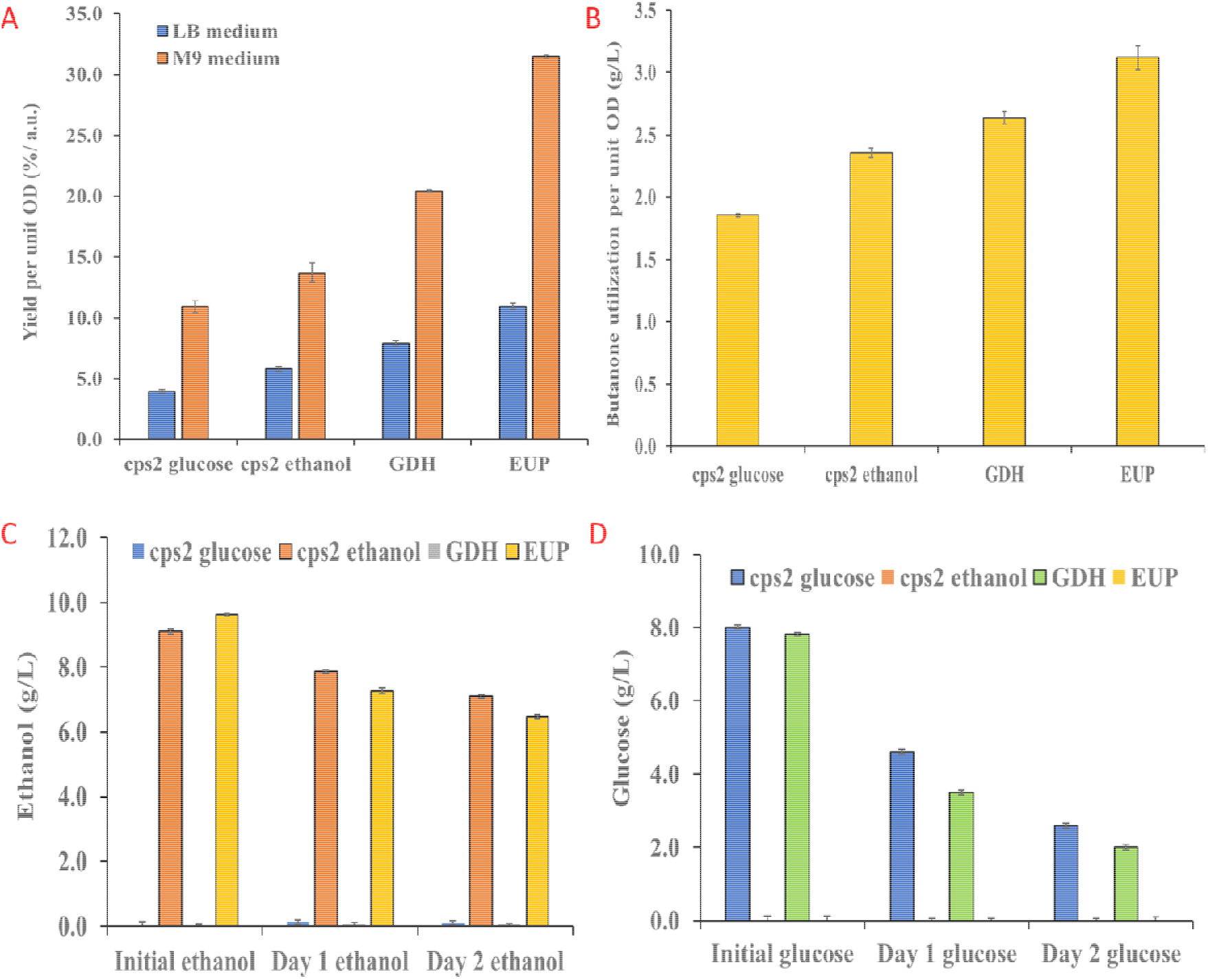
Comparison of biotransformation efficiency of EUP and GDH in resting cells with either 10 g/L glucose or 10 g/L ethanol as co-substrates at 37 °C and 225 rpm under anaerobic conditions. A) EUP delivered higher biotransformation efficiency compared to GDH irrespective of the growth medium in which the resting cells were precultivated, B) Butanone utilisation per unit OD showed the same trend as yield per unit OD across the different categories tested in resting cells precultivated in M9 + 10 g/L ethanol medium, which suggests that amount of NADH regenerated drives butanone utilisation, C) Cells with EUP pathway utilized more ethanol compared to cells with endogenous ethanol utilisation system (*E. coli* cps2), which indicates that heterologous EUP was expressed and was functional, D) More glucose was consumed by cells with GDH compared to cells with endogenous glucose utilisation system (*E. coli* cps2). Inability to fully utilise glucose over 2 days meant that a fundamental limit may exist in glucose utilisation under anaerobic conditions which precludes loading of more glucose into the biotransformation system to enable equimolar comparisons between the two substrates. Data for C and D were obtained from resting cells precultivated in M9 + 10 g/L ethanol medium.

To compare the relative efficiency of two NADH regeneration systems require fair comparison of their substrate utilisation. This then require equal amount of glucose or ethanol to be provided to the cells under the molar basis. But data from Figure 4c and Figure 4d concerning glucose and ethanol utilisation imply that the above idea may not be possible in this system. Specifically, glucose could not be completely consumed over 2 days, which suggests a fundamental limit to glucose consumption may exist in the system. Such a situation then precludes the use of equimolar concentration of glucose and ethanol for determining the relative efficiency of GDH and EUP cofactor regeneration system. The data collected suggests that ethanol utilisation with EUP may be a better NADH regeneration system compared to glucose utilisation with GDH after considering substrate consumption by cells in a whole cell biocatalytic system. Overall, the data and analysis reported here points to the need to consider limitations in substrate uptake by cells during comparison of relative efficiency of different cofactor regeneration systems in whole cell biocatalysis systems.

### Growing cells deliver a driving force that improves biotransformation efficiency over resting cells

Growing cells have been used in many areas of metabolic engineering to drive flux down a designated metabolic pathway to deliver higher product yield. This approach relies heavily on engineering particular nodes or intermediates to be coupled to cell growth, which would subsequently drive flux to move towards the engineered nodes in the metabolic network. In this project, the EUP pathway ends at acetyl-CoA, which is a natural growth promoting metabolite; thus, cell growth should be able to draw flux from acetyl-CoA for building biomass, and hence, help move flux from ethanol to acetyl-CoA that helps regenerate more NADH in support of biotransformation.

Experiment data verified theoretical predictions that growing cell should deliver a higher biotransformation efficiency compared to resting cells (Figure 5a). But, what is of greater interest is the confirmation, from metabolite utilisation and production data, that indeed, growing cells delivered a driving force to improve carbon flux from ethanol to acetyl-CoA and help improve NADH regeneration in support of biotransformation. To do this, both ethanol utilisation and acetate formation was monitored by HPLC. In particular, ethanol utilisation per unit OD revealed that growing cells helped improve ethanol consumption compared to resting cells (Figure 5b). This result validates that growing cells could improve ethanol utilisation by forcing flux to move down the EUP pathway. At the other end of the pathway, observations of increased acetate formation per unit OD in growing cells compared to resting cells again validated that cell growth drove flux from ethanol to acetyl-CoA which is converted to acetate through acetyl-phosphate (Figure 5c). Part of the reason there was high acetate concentration in growing cells despite channelling of acetyl-CoA flux to cell growth in growing cell biotransformation could be the limited cell growth in M9 + 10 g/L ethanol medium (Figure 5d). Such limited growth is opposite to observed good growth of *E. coli* ethcps2 in M9 + 10 g/L ethanol medium under anaerobic conditions without butanone input. This thus indicates that butanone exerted a toxicity effect on *E. coli*, especially with ethanol as a growth substrate.

**Figure 5.**
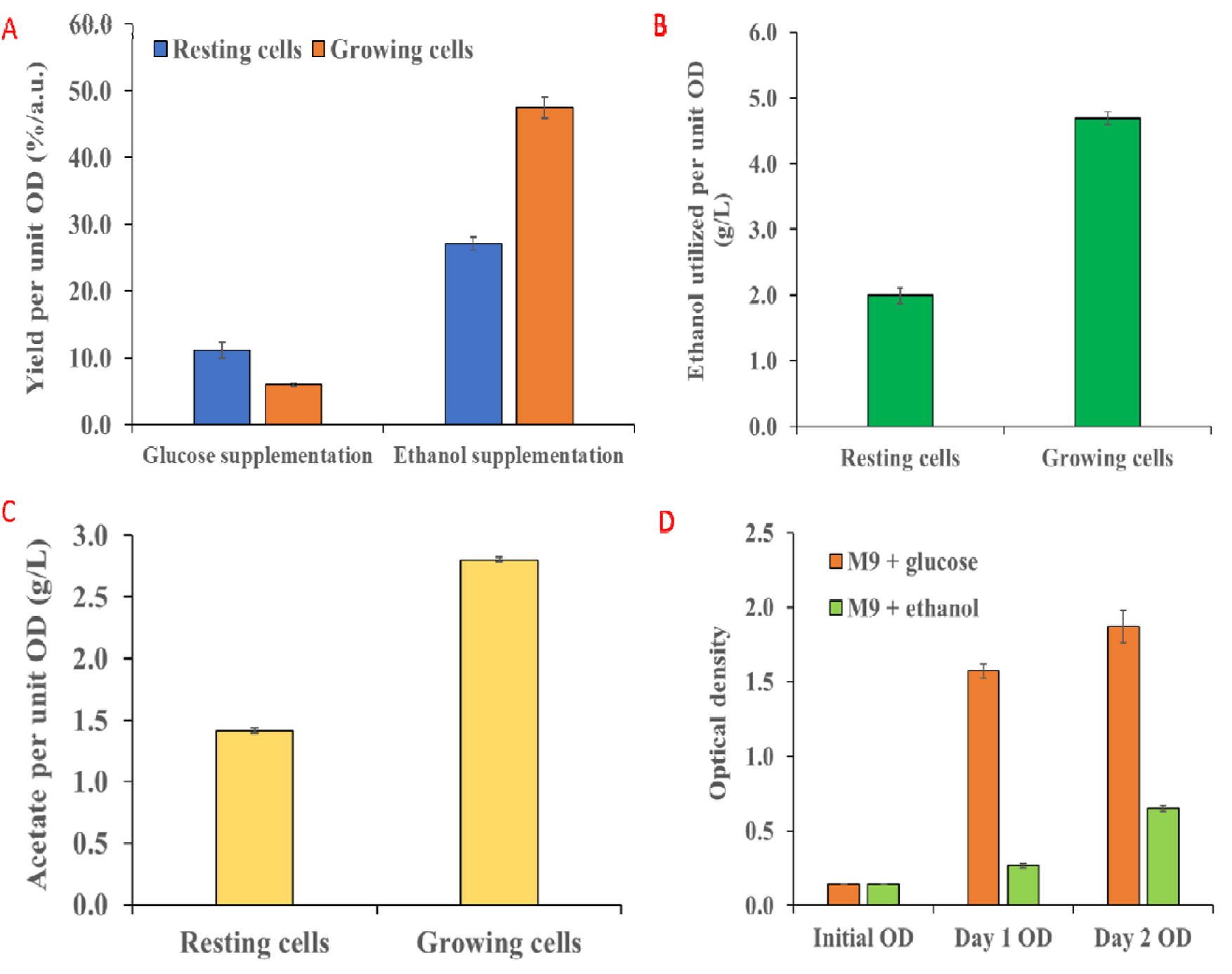
Comparison between the biotransformation efficiency achievable for resting and growing cells either precultivated or grown in M9 + 10 g/L ethanol medium. A) Growing cells delivered a higher biotransformation efficiency over resting cells for those fed with ethanol, B) Ethanol utilisation per unit OD was also higher for growing cells than resting cells, bearing testament to the effect of driving force from cell growth that led to increased ethanol consumption and NADH regeneration, C) Acetate formation per unit OD was also higher for growing cells compared to resting cells, which confirms that growing cells drove greater flux from ethanol to acetyl-CoA and on to acetate through acetyl-phosphate, D) Cells could grow in M9 + 10 g/L ethanol medium during growing cell biotransformation, but growth was retarded probably due to toxicity of butanone. All experiments conducted in this series were performed under anaerobic conditions at 37 °C and 225 rpm.

### Effect of growth medium on biotransformation efficiency in resting and growing cells

Growth medium influences the gene expression pattern of cells, and this may have an impact on biotransformation efficiency such as that manifested by lower expression of NADH dependent enzymes that would compete for NADH originally slated for biotransformation. In the context of this work, the growth medium effect could be present in both resting cell and growing cell biotransformation. This last segment of the work aims to probe how two common growth media (LB and M9 + 10 g/L ethanol) in biocatalysis would influence biotransformation efficiency of butanone in resting and growing cells. Experimental results revealed that in resting cell biotransformation, cells precultivated in M9 + 10 g/L ethanol medium delivered higher biotransformation efficiency compared to those precultivated in LB medium (Figure 6a). The effect was independent of whether glucose or ethanol was used as the co-substrate in resting cell biotransformation, and suggests that observed phenomenon cannot be solely attributed to “training effect” of cells precultivated in medium with ethanol supplementation. In growing cells biotransformation, cells grown in M9 + 10 g/L ethanol medium also delivered higher biotransformation efficiency compared to those grown in LB medium (Figure 6b). Overall, growth medium effect studies indicated that M9 + 10 g/L ethanol medium may be better for improving butanone biotransformation efficiency through enhancing NADH regeneration. A possible reason that could account for the observation could be lower expression of NADH dependent enzymes in cells precultivated or cultivated in M9 + 10 g/L ethanol medium. This way, there would be less competition for regenerated NADH, and more of it could be used by cpsADH to drive butanone conversion into 2-butanol.

**Figure 6.**
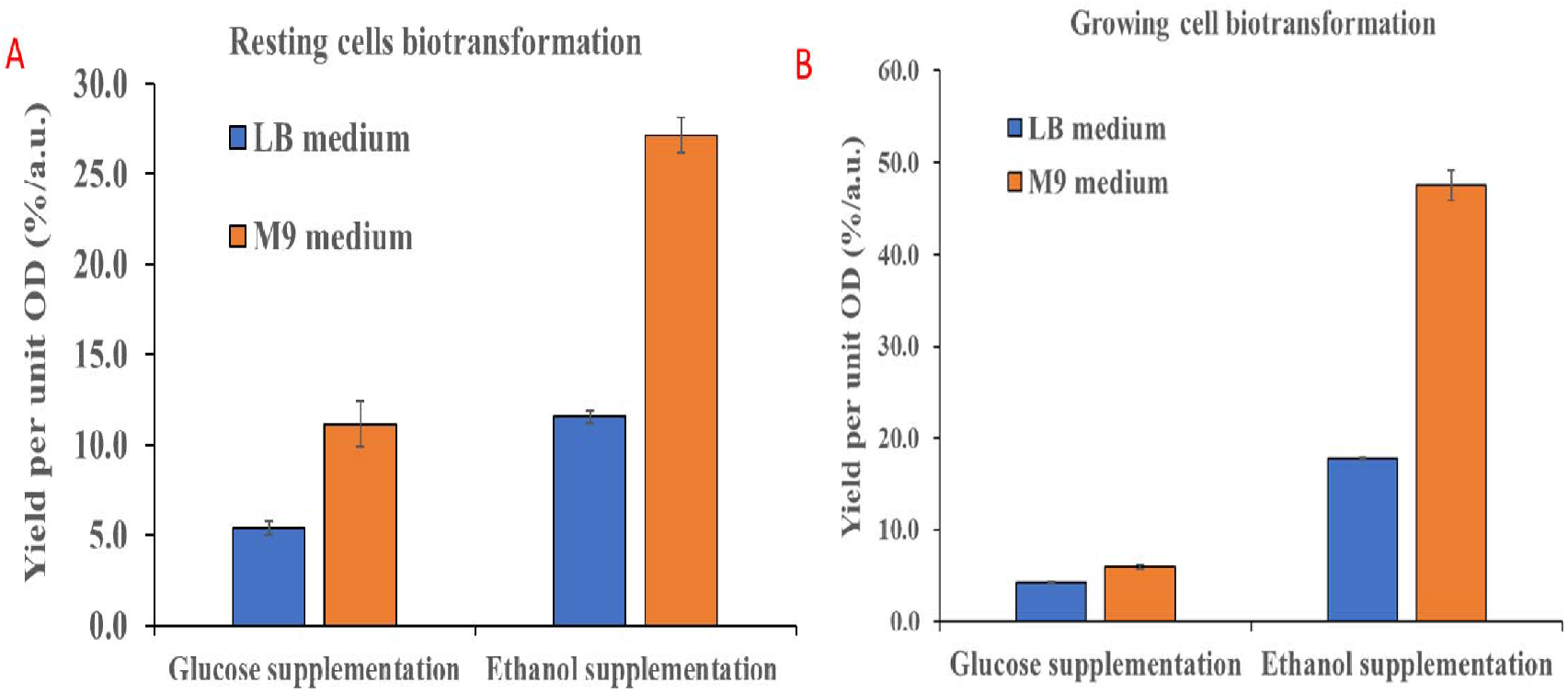
Effect of growth medium on biotransformation efficiency in resting and growing cells. A) Cells precultivated in M9 + 10 g/L ethanol medium delivered higher biotransformation efficiency compared to those precultivated in LB medium during resting cell biotransformation, B) Similarly, cells cultivated in M9 + 10 g/L ethanol delivered higher yield per unit OD compared to those cultivated in LB medium during growing cell biotransformation. All experiments were conducted under anaerobic conditions at 37 °C and 225 rpm.

## Conclusions

Cofactor regeneration is an established technology in biocatalysis, but the search remains for new sacrificial substrate and enzymes capable of delivering enhanced cofactor regeneration at a cheaper cost. In addition, the trend has been moving towards pathway-based cofactor regeneration where the cofactors could be usefully tapped for balancing the redox environment in metabolic engineering application. This work sought to validate theoretical predictions of the utility of a two gene/enzyme ethanol utilisation pathway (EUP) in supplying NADH for a proven ketone to alcohol biotransformation reaction mediated by cpsADH enzyme from *Candida parapsilosis*. Microbial chassis to demonstrate the platform technology is *Escherichia coli* BL21 (DE3).

Experimental results confirm that EUP could be readily integrated with cpsADH to constitute a whole-cell biocatalytic system able to convert ketone (acetophenone or butanone) to alcohol (1-phenylethanol or 2-butanol) in either two-phase or single-phase biotransformation systems. In acetophenone biotransformation, the whole-cell biocatalytic system could tolerate elevated concentration of acetophenone with no evident substrate toxicity effect given the use of an organic phase to solubilize most of the hydrophobic substrate. On the other hand, single phase butanone biotransformation saw evidence of substrate toxicity effect negatively affecting product yield and biotransformation efficiency at elevated substrate concentration. However, EUP pathway could still mediate significant cofactor regeneration and biotransformation under impact from substrate toxicity.

Comparative assessment of EUP and glucose dehydrogenase (GDH) system in butanone biotransformation saw enhanced biotransformation efficiency from EUP compared to GDH in whole cells provided with equal mass concentration of glucose and ethanol. Inability of whole-cell system to fully utilize glucose nevertheless precluded a more comprehensive assessment of the two NADH regeneration systems at an equimolar basis. On the other hand, efforts to address the equilibrium limited product profile from a reversible pathway saw success where higher ethanol concentration drove higher NADH regeneration and better biotransformation efficiency. But, the most prominent characteristic of the EUP pathway remains its ability to be tapped by cell growth process to generate a driving force through the pathway that enhance NADH regeneration. Data supported this theoretical prediction and offered an approach for which EUP pathway could be usefully applied in biocatalysis and metabolic engineering. Finally, evaluation of growth medium influence on biotransformation efficiency saw enhanced yield per unit optical density from cells precultivated or grown in M9 + 10 g/L ethanol over LB medium. The effect is independent of resting or growing cells, and could be due to lower expression of NADH dependent enzymes in cells exposed to M9 + 10 g/L ethanol that reduced competition of NADH originally slated for biotransformation. Observations of enhanced biotransformation from cells precultivated in M9 + 10 g/L ethanol and which were fed with glucose during biotransformation confirmed that the medium effect is not simply due to “training” of cells to more efficiently use ethanol. Overall, demonstrations of the EUP pathway’s capability at supporting both single phase and two-phase biotransformation across two substrates provided another potential useful tool at cofactor regeneration for the biocatalysis and metabolic engineering community. However, constrains in ethanol utilisation highlight opportunities for further engineering the system to afford even higher biotransformation efficiency.

---

**Supplementary Table 1:**
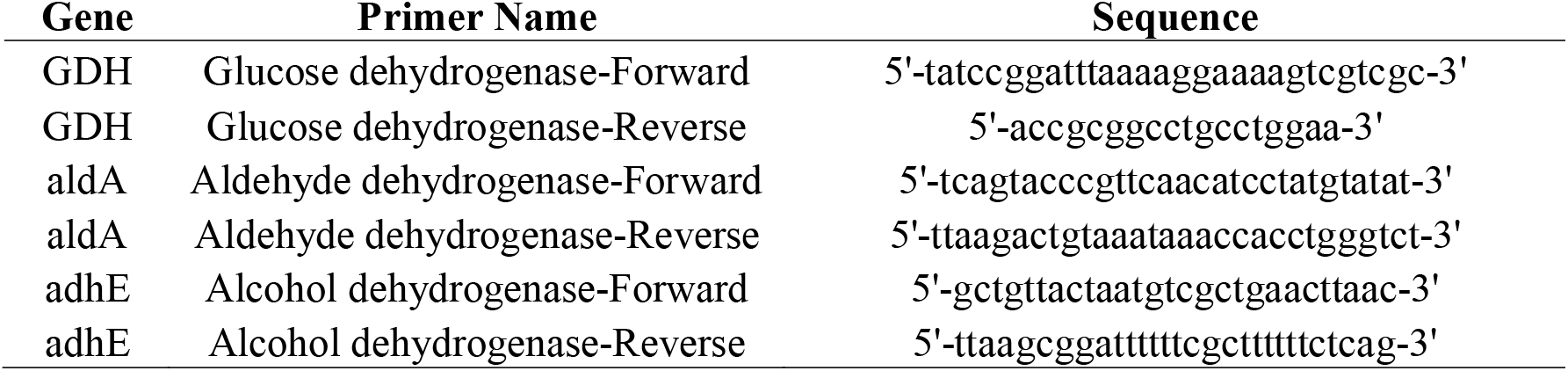
Primers used in the ethanol cofactor regeneration project.

## References

1 Kim, P.-Y., Pollard, D. J. & Woodley, J. M. Substrate Supply for Effective Biocatalysis. Biotechnology Progress 23, 74–82, doi:10.1021/bp060314b (2007).

2 Lin, B. & Tao, Y. Whole-cell biocatalysts by design. Microbial Cell Factories 16, 106, doi:10.1186/s12934-017-0724-7 (2017).

3 Xiao, Z. et al. A Novel Whole-Cell Biocatalyst with NAD+ Regeneration for Production of Chiral Chemicals. PLOS ONE 5, e8860, doi:10.1371/journal.pone.0008860 (2010).

4 Wachtmeister, J. & Rother, D. Recent advances in whole cell biocatalysis techniques bridging from investigative to industrial scale. Current Opinion in Biotechnology 42, 169–177, doi:https://doi.org/10.1016/j.copbio.2016.05.005 (2016).

5 Shah, S., Sunder, A. V., Singh, P. & Wangikar, P. P. Characterization and Application of a Robust Glucose Dehydrogenase from Paenibacillus pini for Cofactor Regeneration in Biocatalysis. Indian Journal of Microbiology 60, 87–95, doi:10.1007/s12088-019-00834-w (2020).

6 Schüürmann, J., Quehl, P., Lindhorst, F., Lang, K. & Jose, J. Autodisplay of glucose-6-phosphate dehydrogenase for redox cofactor regeneration at the cell surface. Biotechnology and Bioengineering 114, 1658–1669, doi:10.1002/bit.26308 (2017).

7 Mourelle-Insua, Á., Aalbers, F. S., Lavandera, I., Gotor-Fernández, V. & Fraaije, M. W. What to sacrifice? Fusions of cofactor regenerating enzymes with Baeyer-Villiger monooxygenases and alcohol dehydrogenases for self-sufficient redox biocatalysis. Tetrahedron 75, 1832–1839, doi:https://doi.org/10.1016/j.tet.2019.02.015 (2019).

8 Qian, W.-Z. et al. Evolution of Glucose Dehydrogenase for Cofactor Regeneration in Bioredox Processes with Denaturing Agents. ChemBioChem n/a, doi:10.1002/cbic.202000196 (2020).

9 Zhang, Y., Wang, Y., Wang, S. & Fang, B. Engineering bi-functional enzyme complex of formate dehydrogenase and leucine dehydrogenase by peptide linker mediated fusion for accelerating cofactor regeneration. Engineering in Life Sciences 17, 989–996, doi:10.1002/elsc.201600232 (2017).

10 Shaked, Z. e. & Whitesides, G. M. Enzyme-catalyzed organic synthesis: NADH regeneration by using formate dehydrogenase. Journal of the American Chemical Society 102, 7104–7105, doi:10.1021/ja00543a038 (1980).

11 Tishkov, V. I. & Popov, V. O. Catalytic mechanism and application of formate dehydrogenase. Biochemistry (Moscow) 69, 1252, doi:10.1007/s10541-005-0071-x (2004).

12 Rocha-Martín, J. et al. New biotechnological perspectives of a NADH oxidase variant from Thermus thermophilus HB27 as NAD+-recycling enzyme. BMC Biotechnology 11, 101, doi:10.1186/1472-6750-11-101 (2011).

13 Geueke, B., Riebel, B. & Hummel, W. NADH oxidase from Lactobacillus brevis: a new catalyst for the regeneration of NAD. Enzyme and Microbial Technology 32, 205–211, doi:https://doi.org/10.1016/S0141-0229(02)00290-9 (2003).

14 Matsumoto, J., Higuchi, M., Shimada, M., Yamamoto, Y. & Kamio, Y. Molecular Cloning and Sequence Analysis of the Gene Encoding the H2O-forming NADH Oxidase from Streptococcus mutans. Bioscience, Biotechnology, and Biochemistry 60, 39–43, doi:10.1271/bbb.60.39 (1996).

15 Pongtharangkul, T. et al. Kinetic properties and stability of glucose dehydrogenase from Bacillus amyloliquefaciens SB5 and its potential for cofactor regeneration. AMB Express 5, 68, doi:10.1186/s13568-015-0157-9 (2015).

16 Riebel, B. R., Gibbs, P. R., Wellborn, W. B. & Bommarius, A. S. Cofactor Regeneration of NAD+ from NADH: Novel Water-Forming NADH Oxidases. Advanced Synthesis & Catalysis 344, 1156–1168, doi:10.1002/1615-4169(200212)344:10<1156::AID-ADSC1156>3.0.CO;2-# (2002).

17 Liang, H. et al. Constructing an ethanol utilization pathway in Escherichia coli to produce acetyl-CoA derived compounds. bioRxiv, 2020.2004.2014.041889, doi:10.1101/2020.04.14.041889 (2020).

18 Yamamoto, H., Kawada, N., Matsuyama, A. & Kobayashi, Y. Cloning and Expression in Escherichia coli of a Gene Coding for a Secondary Alcohol Dehydrogenase from Candida parapsilosis. Bioscience, Biotechnology, and Biochemistry 63, 1051–1055, doi:10.1271/bbb.63.1051 (1999).

19 Ma, X. et al. A standard for near-scarless plasmid construction using reusable DNA parts. Nature Communications 10, 3294, doi:10.1038/s41467-019-11263-0 (2019).

