## Supplementary material for "Developing an ethanol utilisation pathway based NADH regeneration system in *Escherichia coli*"

**Supplementary materials of “Developing an ethanol utilization pathway based NADH regeneration system in *Escherichia coli*”**

Wenfa Ng

Department of Chemical and Biomolecular Engineering, National University of Singapore,

**Supplementary figures**

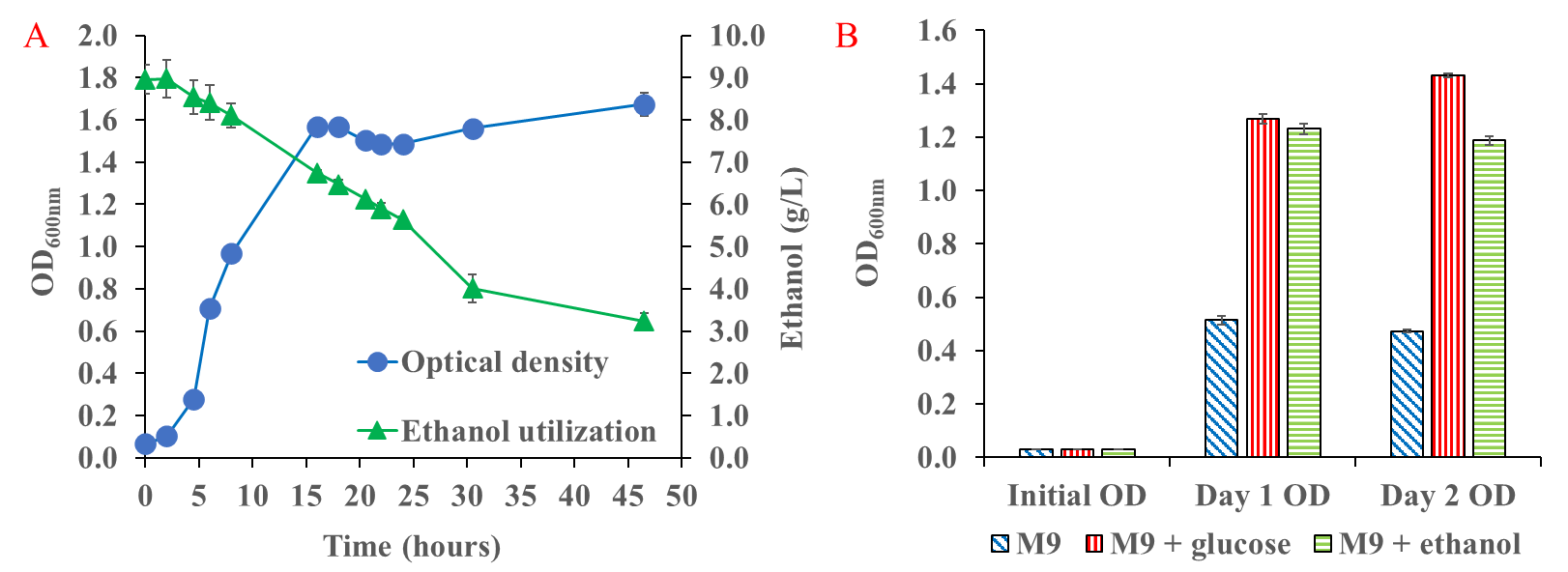

**Fig. S1.** EUP could deliver growth to *E. coli* under aerobic and anaerobic conditions. A. Growth of *E. coli* ethcps2 in M9 + 10 g/L ethanol medium under aerobic conditions at 37 ^o^C and 225 rpm in 125 mL shake flasks, B. Growth of *E. coli* ethcps2 in M9 medium with different carbon source under anaerobic conditions at 37 ^o^C and 225 rpm in 2.0 mL glass HPLC vials. The basal M9 medium contains 1 g/L yeast extract.

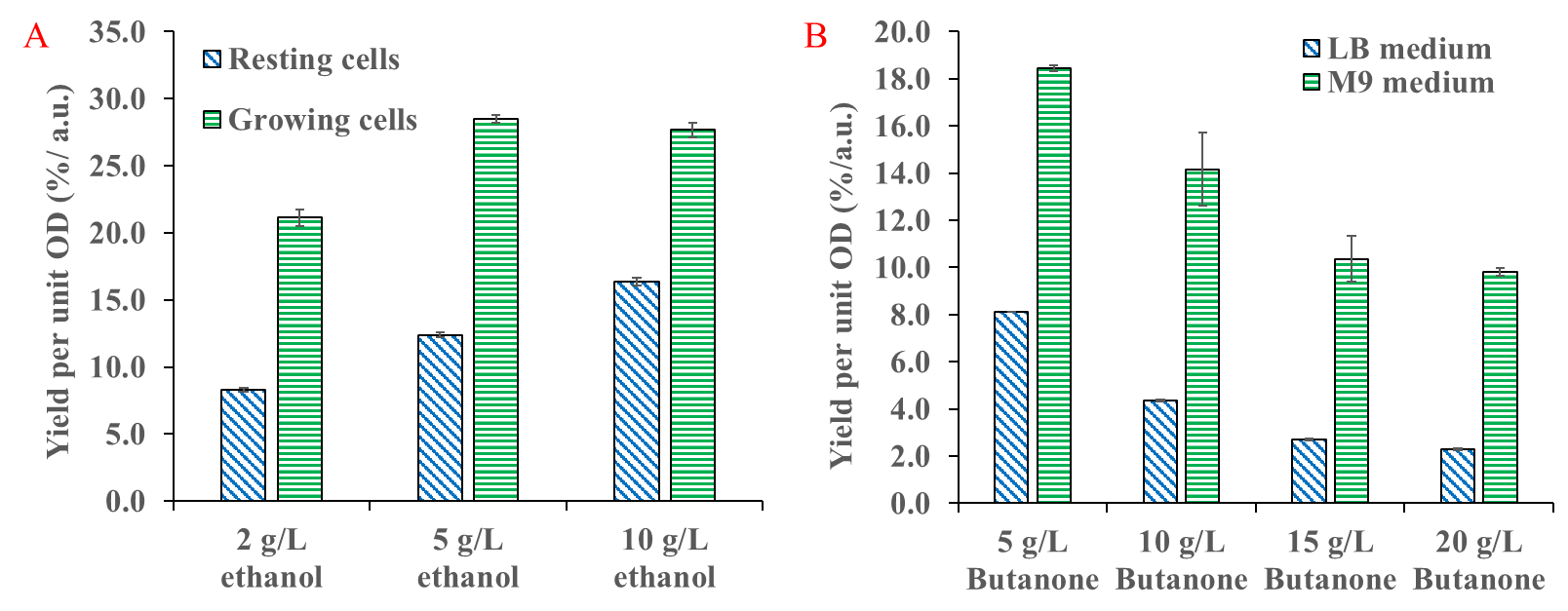

**Fig. S2.** Effect of ethanol and butanone concentration on butanone biotransformation efficiency in *E. coli* ethcps2. A. Biotransformation efficiency positively correlated with ethanol concentration in resting cells, but toxicity effect was detected at 10 g/L ethanol in growing cell biotransformation, B. Increase in butanone concentration resulted in decrease in biotransformation efficiency for resting cells precultivated in either LB or M9 + 10 g/L ethanol medium. Results indicated existence of substrate toxicity effect in single phase butanone biotransformation. All experiments were performed at 37 ^o^C and 225 rpm under anaerobic conditions.

| **Supplementary Table 1: Primers used in the ethanol cofactor regeneration project** | | |
| --- | --- | --- |
| **Gene** | **Primer Name** | **Sequence** |
| adh2 | Alcohol dehydrogenase-Forward | 5'-tctattccagaaactcaaaaagccattatctt-3' |
| adh2 | Alcohol dehydrogenase-Reverse | 5'-tttagaagtgtcaacaacgtatctaccagc-3' |
| ada | Acetaldehyde dehydrogenase-Forward | 5'-gaacactccgtgatcgaacc-3' |
| ada | Acetaldehyde dehydrogenase-Reverse | 5'-ggcaatacggaacatatccaccag-3' |
| cpsADH | Alcohol dehydrogenase-Forward | 5'-tcaatcccgtcgtcccaata-3' |
| cpsADH | Alcohol dehydrogenase-Reverse | 5'-cgggttaaagacgacgcg-3' |
| GDH | Glucose dehydrogenase-Forward | 5'-tatccggatttaaaaggaaaagtcgtcgc-3' |
| GDH | Glucose dehydrogenase-Reverse | 5'-accgcggcctgcctggaa-3' |

| **Supplementary Table 2: Plasmids used in the project** | | |
| --- | --- | --- |
| **Plasmid name** | **Promoter** | **Genes** |
| p76 | pthrC3 | cpsADH alcohol dehydrogenase |
| eth1 | T7 | adh2 (alcohol dehydrogenase)  and ada (acetaldehyde dehydrogenase) |
| eth2 | pthrC3 | adh2 (alcohol dehydrogenase)  and ada (acetaldehyde dehydrogenase) |
| GDH1 | pthrC3 | GDH glucose dehydrogenase |

| **Supplementary Table 3: Bacterial strains used in the project** | | |
| --- | --- | --- |
| **Strain name** | **Host strain** | **Plasmids carried by strain** |
| *E. coli* cps2 | *E. coli* BL21 (DE3) | p76 plasmid carrying cpsADH under pthrC3 control |
| *E. coli* ethcps1 | *E. coli* BL21 (DE3) | eth1 plasmid carrying EUP under T7 promoter,  and p76 plasmid with cpsADH under pthrC3 control |
| *E. coli* ethcps2 | *E. coli* BL21 (DE3) | eth2 plasmid carrying EUP under pthrC3 promoter,  and p76 plasmid with cpsADH under pthrC3 control |
| *E. coli* GDHcps1 | *E. coli* BL21 (DE3) | GDH1 plasmid carrying GDH under pthrC3 promoter,  and p76 plasmid with cpsADH under pthrC3 control |

**Conflicts of interest**

The author declares no conflicts of interest.

**Funding**

The author thank the National University of Singapore for financial support.
